# Material Damage to Multielectrode Arrays after Electrolytic Lesioning is in the Noise

**DOI:** 10.1101/2025.03.26.645429

**Authors:** Alice Tor, Stephen E Clarke, Iliana E Bray, Paul Nuyujukian, Brain Interfacing Laboratory

## Abstract

The quality of stable long-term recordings from chronically implanted electrode arrays is essential for experimental neuroscience and brain-computer interfaces. This work uses scanning electron microscopy (SEM) to image and analyze eight 96-channel Utah arrays previously implanted in motor cortical regions of four subjects (subject H = 2242 days implanted, F = 1875, U = 2680, C = 594), providing important contributions to a growing body of long-term implant research leveraging this imaging technology. Four of these arrays have been used in electrolytic lesioning experiments (H = 10 lesions, F = 1, U = 4, C = 1), a novel electrolytic perturbation technique using small direct currents. In addition to surveying physical damage, such as biological debris and material deterioration, this work also analyzes whether electrolytic lesioning created damage beyond what is typical for these arrays. These findings also indicate that there are no statistically significant differences between the damage observed on normal electrodes versus electrodes used for electrolytic lesioning, providing evidence that electrolytic lesioning does not significantly affect the quality of chronically implanted electrode arrays. Finally, this work also includes the largest collection of single-electrode SEM images for previously implanted multielectrode Utah arrays, spanning eleven different intact arrays and one broken array. As the clinical relevance of chronically implanted electrodes with single-neuron resolution continues to grow, these images may be used to provide the foundation for a larger public database and inform further electrode design and analyses.

## 1 Introduction

In the past two decades, the advancement of implanted electrode arrays has drastically shifted the field of neuroscience from theories based on individually recorded neurons to those based on the systems-level activity of neuron populations [1–3]. Specifically, the 96-channel Utah array (Blackrock Neurotech, Salt Lake City, UT) has been used to make pivotal discoveries in motor systems neuroscience and develop clinically impactful brain-computer interfaces [2, 4–8]. These electrode arrays are widely used in neuroprosthetics research for patients with motor disorders and have been used for a variety of purposes, such as movement and speech decoding [9, 10]. Much of this work in humans relies on the translation of results discovered in non-human primates (NHPs), a crucial animal model in neuroscience research [11, 12]. One useful application of these arrays in animal studies is microstimulation and electrolytic lesion experiments [13]. Electrolytic lesioning results in small, controllable, and precise amounts of neuron loss. This makes it a helpful technique to study damage to targeted areas of the brain, serving as a causal tool for basic scientific inquiry and as a model for clinical injury-based neuron loss events at a small scale.

Electrolytic lesioning has been experimented with since the 19th century [14], and methods have been researched in many species, including rodents, cats, dogs, monkeys, and humans [15–17]. In electrolytic lesion experiments, researchers pass current through implanted electrodes in order to mark the position of electrodes or create lesions in brain tissue [17, 18]. These electrolytic lesions can be created using one electrode (unipolar) or by using two electrodes (bipolar) [16, 19]. In particular, recent work has established a novel lesioning technique to create consistent, controllable electrolytic lesions in NHPs using already-implanted Utah arrays, without damaging the ability to record from the array [20]. However, while recording capabilities seem to remain stable, the physical effect of electrolytic lesioning on the recording array needs further characterization.

Previous studies have shown temporally decreasing performance of collected signals from long-term implanted arrays, thought to be a consequence of the immune response, mechanical movement of the array relative to surrounding tissue, and physical degradation of the array [21–23]. This is also reflected in studies that specifically examine the damages that occur to electrode arrays while they are implanted in NHPs [24, 25]. In particular, the use of scanning electrode microscopy (SEM) has been used to visually examine and analyze the effect of neural implantation on arrays with extremely high definition [25–29]. These studies, which span both NHPs and humans, provide direct evidence that the observable physical damage incurred by implanted arrays has direct impact on the array’s performance.

The Utah array is a silicon-based array manufactured with several layers of different materials [30]. Silicon electrodes are first etched out of a glass and silicon base. Each individual electrode is then metallized at the tip. Finally, a layer of parylene C coats all but these metallized tips to help isolate and insulate each electrode [31, 32]. Both *in vitro* and *in vivo* testing of electrode arrays revealed environmental damage to these materials, such as cracking, textural defects, and degradation in response to the brain’s temperature and salinity [33, 34]. The immune response of the brain also damages the electrodes due to effects like glial scarring (gliosis) and inflammation [34–36]. This damage may be exacerbated by the surgical techniques used during implantation, which include pushing the electrode array into cortex and tethering the implant to the skull [35, 37, 38]. These results are confirmed in the previously mentioned SEM analyses [25–29].

Additionally, the Utah array is available with both platinum and iridium oxide-based coatings. Aggressive electrical stimulation is known to dissolve platinum-based electrodes [39, 40]. Other studies have shown iridium oxide to be more resistant to stimulation-related damage, but not completely insusceptible [26, 41–43]. However, the physical electrode damage associated with electrolytic lesioning remains to be studied.

This work comprises images of eleven different multielectrode Utah arrays (ten 96-channel, one 64-channel). Of these imaged arrays, eight arrays chronically implanted in regions of the motor cortex for varying amounts of time were analyzed further (Monkey H = 2242 days, F = 1875, U = 2680, C = 594). Four out of eight of these arrays were used to perform electrolytic lesioning experiments (n = 4 monkeys; H = 10 lesions, F = 1, U = 4, C = 1). Additionally, the image set includes images of a broken 96-channel Utah array, which was shattered during explant but remained intact while implanted. This set of images represents the largest publicly available collection of high-quality, single-electrode SEM images of explanted multielectrode Utah arrays. A total of 938 individual electrodes are available to view, along with 11 additional images from the broken array. Specific details for each imaged array are available in Supplemental Table 1.

## 2 Methods

### Electrolytic Lesioning

All lesions were performed following previously described procedures[20]. To summarize, each lesion is created by passing direct current (DC) between two electrodes on the array. Thus, each lesion is bipolar, not unipolar, and is produced without a pulse/frequency-based protocol. Exact parameters for each lesion are available in Supplemental Table 6.

### Array Preparation

Monkey H, F, and C were each implanted with two 96-channel Utah arrays in M1 (primary motor cortex) and PMd (dorsal premotor cortex). H’s M1 array was used for electrolytic lesioning experiments. F’s and C’s PMd arrays were used for electrolytic lesioning experiments. Monkey U was implanted with three 96-channel arrays; two in M1, and one in PMd. U’s lateral M1 array was used for electrolytic lesioning experiments. U’s medial M1 array was not used for recording, and was severely damaged during extraction. Monkey H’s arrays were implanted for 2242 days (electrolytic lesions on days 2088, 2129, 2136, 2164, 2172, 2180, 2187, 2192, 2221, 2228). Monkey F’s arrays were implanted for 1875 days (electrolytic lesion on day 1875). Monkey U’s arrays were implanted for 2680 days (electrolytic lesions on days 1215, 1250, 1264, 1286). Monkey C’s arrays were implanted for 594 days (electrolytic lesion on day 304). To produce each electrolytic lesion, current was passed between two electrodes on the array. Further details on each monkey’s arrays, as well as any additional imaged arrays, are available in Supplemental Table 1. Further details on lesioning procedures are available in Bray et al., 2024 [20].

All arrays used in this work were multielectrode Utah arrays (Blackrock Neurotech, Salt Lake City, UT). Monkey C was implanted with an array with electrode shank lengths of 1.5 mm, while Monkeys H, F, and U were implanted with arrays with electrode shank lengths of 1 mm. Monkey U and C’s arrays were manufactured with an iridium oxide metal coating, while H and F’s arrays were manufactured with platinum coatings.

Arrays were first soaked in either a 0.9% saline or 10% formalin solution immediately following explant surgery, then stored separately in either water, formalin, ethanol, or dry in a closed container. Prior to imaging, all arrays were allowed to dry without any further cleaning or preparation.

### Scanning Electron Microscopy

Scanning Electron Microscopy (SEM) images were collected using a Hitachi TM4000+ Tabletop SEM (Hitachi Ltd., Tokyo, Japan). Arrays were mounted on PELCO SEMClip^TM^ Pin Mounts (Ted Pella, Redding, CA) using conductive copper tape, carbon tape, and included clip mounts. Arrays were not sputter-coated for imaging. Due to the amount of organic material left on the arrays, images were collected in low-vacuum mode to avoid distortion from accumulated electrical charge of the non-conductive material. Full array images were collected with the arrays flat on the mount surface, while single-electrode images were collected with the arrays positioned at an approximate 45*^◦^*angle. Most images were collected using a mixed backscatter/secondary electron (BSE/SE) mode with a low accelerating voltage of 5kV, although modes and voltage values were adjusted for certain electrodes to optimize image clarity. Specific settings for each collected image are included in Figure 1.

**Figure 1:** Interactive Display: Video demonstrates how to use the interactive display of all captured SEM array images, available through the provided link.Once an array has been selected, a diagram of each array’s anatomical position (derived from surgical drawings and notes) appears, along with an SEM image of the array. All array images are displayed with the wire bundle to the right side and with electrode tips facing the viewer. Specific electrodes may be selected from the array image; SEM images of each electrode and their scores in each of the five damage categories will appear along with any additional notes. Furthermore, examples of specific scores for each damage type are selectable through an additional drop-down menu for each array. Similarly, electrodes used for electrolytic lesioning are also selectable through an additional drop-down menu for each array.

### Damage Scoring

A prior study categorized damage observable with SEM into six distinct types: abnormal debris, tip breakage, coating cracks (cracking of the metal coating around the silicon core of an electrode), parylene C cracks, parylene C delamination, and shank fracture [27]. Similarly, damage to the arrays in this study were scored in five categories: abnormal debris (AB), silicon tip breakage (TB), platinum coating cracks (CC), parylene C coating cracks (PC), and parylene C coating delamination (PD). Each recording electrode was scored from 0 to 3 in five different categories, as described above. Electrodes with shank fractures (SF) were not scored, as these fractures may have reasonably occurred during explantation and handling of the arrays separate from the damage incurred while in the brain. Each electrode was scored between 0-3, where 0 indicated no visible effects, 1 indicated visible effects but likely no impact on recording ability, 2 indicated possible impact on recording ability, and 3 indicated likely impact on recording ability. Scores were given without prior knowledge of which electrodes had been used for electrolytic lesioning. Although this scoring system is subjective and based on visual observation, all images used to compute these scores are publicly available in Figure 1 for independent evaluation.

### Statistical Tests

First, the Pearson’s correlation coefficient was calculated between each subcategory of damage to determine whether any types of damage were related. To explore potential spatial relationships, this analysis was repeated for each radial ring of electrodes, as shown in Supplemental Figure 4.

Statistical significance was also assessed between electrodes used in the electrolytic lesion experiments versus those which had not. The Shapiro-Wilk test indicated that scores did not follow a normal distribution; therefore, nonparametric methods such as the Mann-Whitney U test and the Levene test were used to compare the underlying statistics of each array’s electrode population. The null hypothesis of the Mann-Whitney U test is that sample means of two sets of scores are the same. The null hypothesis of the Levene test is that the sample variance of two sets of scores are the same.

Similarly, statistical significance was assessed as described above between lesioning versus normal electrodes, which were pooled across all four arrays used for electrolytic lesioning. Finally, this analysis was repeated between electrode scores across pairs of lesioning and non-lesioning arrays implanted in the same subject for the same length of time.

## 3 Results

All collected images, along with their associated scores, are available in interactive Figure 1. Each of the twelve arrays and individual electrodes may be selected and viewed. Representative electrodes for each category and damage score, as well as electrodes used for lesioning, are also viewable for each array in an additional drop-down menu when available. Rough anatomical layouts of the array’s implant location are also included. Electrode numbering maps are available in Supplemental Figure 1.

The first set of analyses specifically considers the four arrays used in electrolytic lesioning experiments. Scores for each damage category, along with total scores, are visualized as heatmaps in Figure 2. Median total scores for each radial section of the array show that the outermost electrodes tend to have the greatest total observed damage across all four arrays. Generally, electrode damage decreases when moving to the center of the array. Shank fractures tend to be spatially close on each array, suggesting that shared mechanical trauma caused these sections of fracture. As mentioned in the Methods section, these fractures may have reasonably occurred during explantation and handling, as no diminished recording performance of these fractured electrodes was observed while the array was actively implanted. Additional heatmaps are available for the four additional arrays implanted in NHP but not used for lesioning in Supplemental Figure 2.

**Figure 2:**
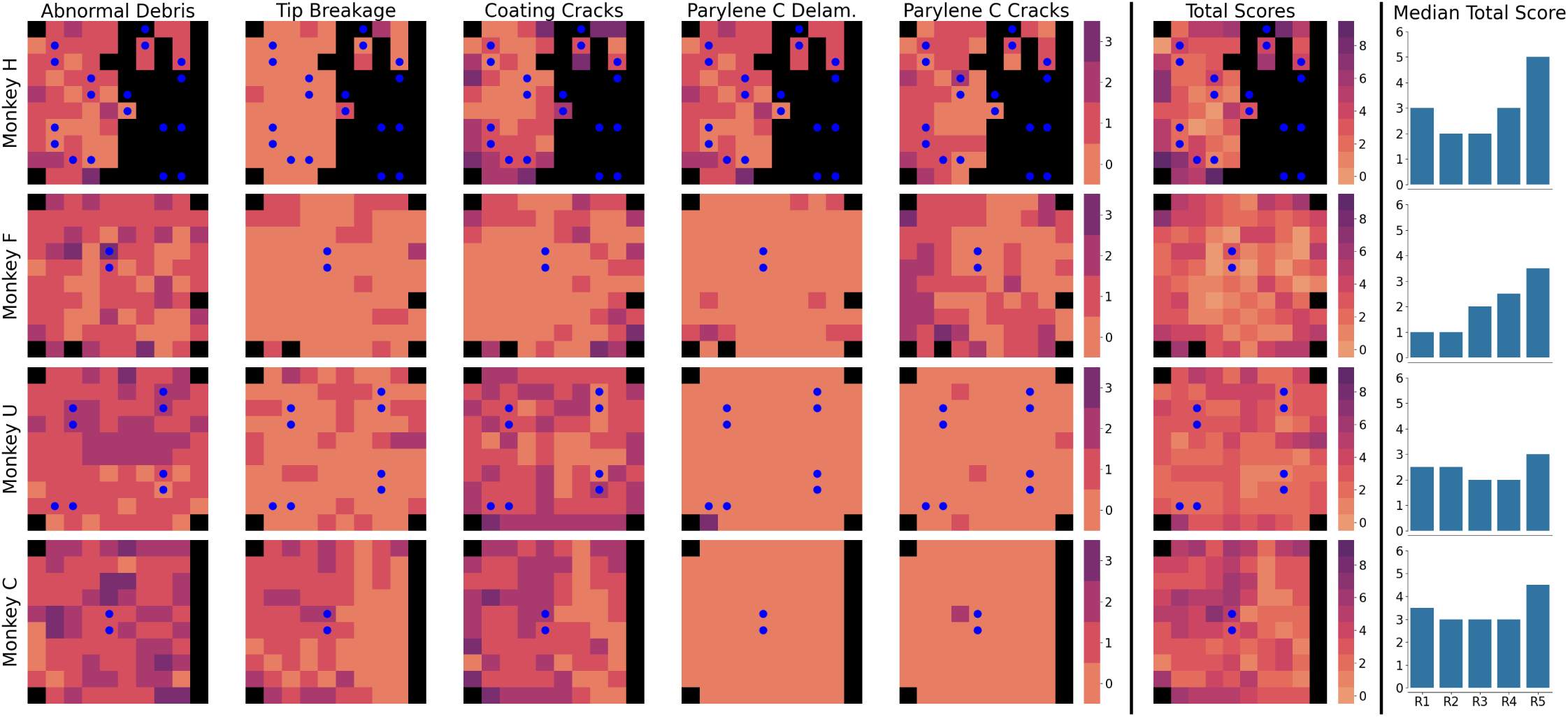
Heatmaps of damage scores (0-3) across the five identified types of damage and across the four imaged lesioning arrays. Electrodes are displayed using the orientation in Figure 1 (electrode tips facing viewer, wire bundle on the right). Second-to-rightmost column displays summed damage scores for each array across the five types of damage. Electrodes used for electrolytic lesioning are denoted with blue dots. Median summed scores for each radial section of the array are plotted in the bar charts to the right of the heatmaps. Ring layout and numbering information are available in Supplementary Figure 1. Unwired electrodes (electrodes not wire bonded at time of manufacture) and electrodes with shank fractures are ignored and displayed in black, as they are not scored.

Histograms of damage scores in each category, along with average score distributions per array, are shown in Figure 3. For each category, histograms of electrodes used for lesioning (black) are stacked on top of histograms of normal electrodes (gray). Additionally, the average distribution of scores is shown for each array. These plots indicate that there is no observable difference in the distribution of scores given to electrodes used for lesioning versus those not. Additional histograms are available for the four additional arrays implanted in NHP but not used for lesioning in Supplemental Figure 3.

**Figure 3:**
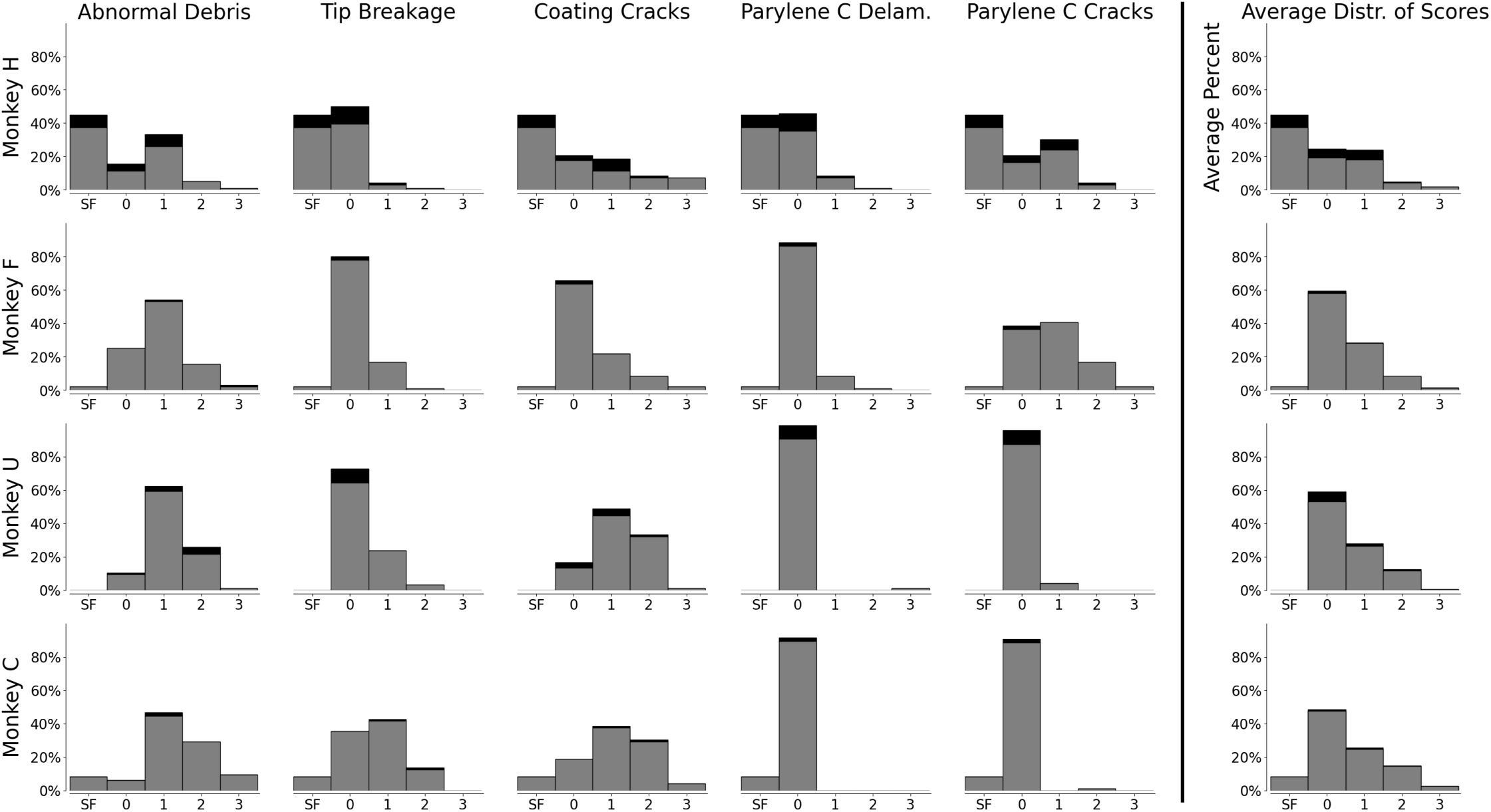
Stacked histograms of damage scores (0-3) across the five identified types of damage and across the four imaged lesioning arrays. Gray indicates normal electrodes, and black indicates lesioning electrodes. The rightmost column displays the average distribution of damage scores across the five types of damage. Electrodes with shank fractures (SF) are ignored, as they are not scored.

The Shapiro-Wilk test for normality returned a significant *p*-value for almost all sets of scores for each array and type of damage. Similarly, the Shapiro-Wilk test for normality returned a significant *p*-value for almost all sets of scores for each array and type of damage when split into populations used and not used for lesioning, but was not performed where electrode count was too small to carry out the test (C and F; lesioning electrodes = 2). In the majority of cases that returned an insignificant *p*-value, certain types of damage were not present on the array and thus scores were uniformly zero (Monkey U’s M1 tip breakage, parylene C delamination, and parylene C cracking scores; Monkey C’s PMd parylene C delamination scores and M1 parylene C cracking scores). In all other cases where the Shapiro-Wilk test returned an insignificant *p*-value (Monkey H’s total damage scores on lesioning electrodes, Monkey U’s coating cracks scores), low electrode counts may reduce the statistical power of the test (H = 18 lesioning electrodes, U = 8). This indicates that scores are largely not normally distributed and that nonparametric statistical tests should be used to evaluate differences between populations. All Shapiro-Wilk test *p*-values are available in Supplemental Table 2.

For each category of damage, as well as the total damage score, no statistically significant difference was identified between lesioning and non-lesioning electrodes on the same array. Similarly, for each category of damage, as well as the total damage score, no statistically significant difference was identified between lesioning and non-lesioning electrodes pooled across all four arrays. This indicates that the electrode populations used for lesioning and those not used for recording only do not have statistically different sample means or variances, suggesting they are samples of the same underlying population. These *p*-values are included in Supplemental Table 3. However, again, it is important to note that these tests done on individual arrays have limited power due to low electrode counts (H = 18 lesioning electrodes, F = 2, U = 8, C = 2).

This next set of analyses compares entire population statistics of all eight NHP arrays. Both the Mann-Whitney U and Levene tests resulted in inconsistently significant/non-significant *p*-values when comparing lesioning arrays against non-lesioning arrays within the same implanted animal. This indicates inconsistency in overall distribution and variance of data. This suggests that there are likely large differences between the amount and intensity of material damage dealt to implanted arrays even when implanted in the same subject. These results do not determine whether or not electrolytic lesioning experiments contribute to the magnitude of these differences, but the inconsistency in the significance of the above results may indicate that other factors, such as the brain’s varied immune response, differences in mechanical forces, and disparities in external handling, may play a large role in damage to these arrays. These p-values are included in Supplemental Table 4. The Pearson correlation matrix across all four arrays for the five different damage types is shown in Figure 4. This plot demonstrates that there is overall low pairwise correlation between each type of damage, with the exception of coating cracks and tip breakage, which display moderate positive correlation (*r* = 0.47, Bonferroni-corrected *p <* 0.05). These results align with with previous comparisons on the distribution of damage scores, which showed that damage score tended to vary across the five different types. Additional correlation plots for each of the five array layout rings are available in Supplemental Materials. Raw *r* and *p*-values for each test are also available in Supplemental Table 5.

**Figure 4:**
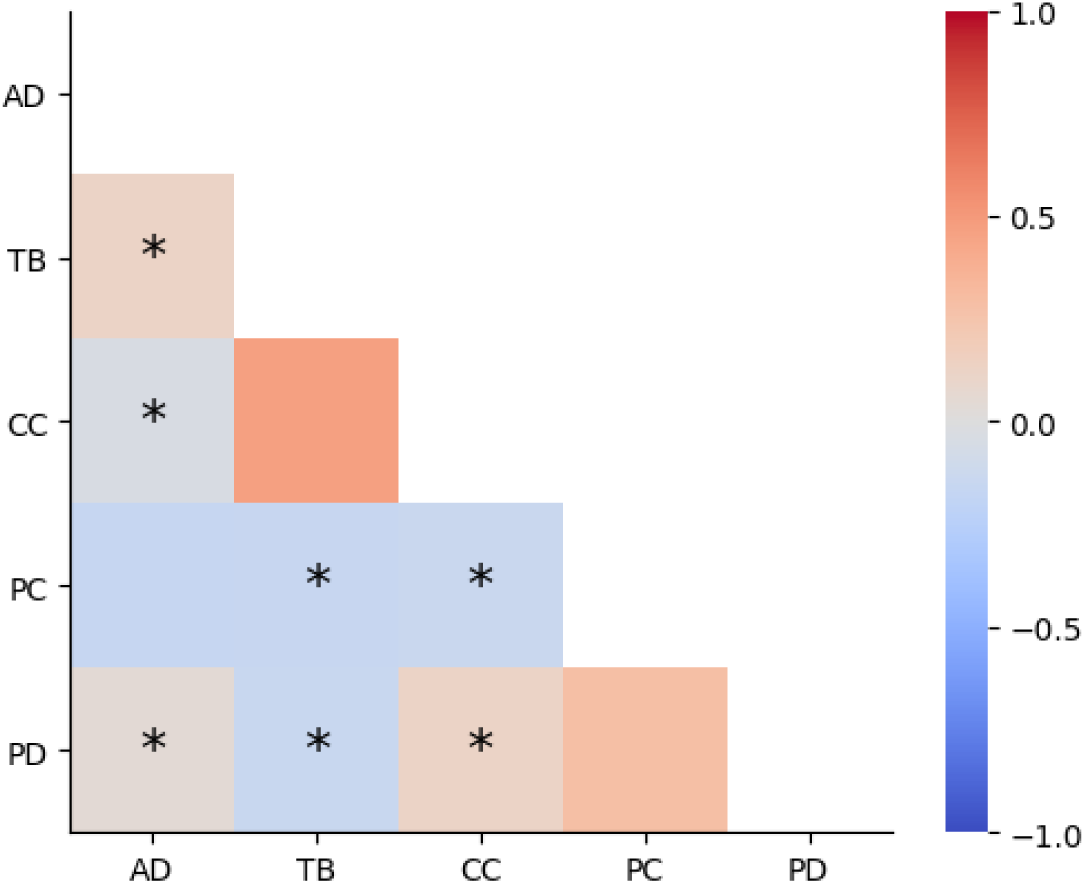
Correlation plot (Pearson’s R) across the five different damage types. AD = abnormal debris, TB = tip breakages, CC = metal coating cracks, PC = parylene C cracks, PD = parylene C delamination. Results demonstrate overall low correlation (magnitude *<* 0.25) across different damage types, with the exception of coating cracks and tip breakage, with a correlation coefficient of 0.47. Test values with Bonferroni-corrected *p <* 0.05 are displayed with asterisks. Raw p-values are separately available in Supplementary Table 2. Electrodes with shank fractures are ignored, as they are not scored.

## 4 Discussion

Overall, these results collectively demonstrate that there is no obvious, statistically significant difference between observed damage to electrodes used versus not used for electrolytic lesioning. This indicates that any material effects potentially caused by electrolytic lesioning are indistinguishable from the typical noise seen in long-term implanted arrays. Because this method of lesioning involves passing weak direct current into the brain over relatively short time periods, its physical effects on the array electrodes are similar to alternating current stimulation. A recent study found that SEM-visible damage caused by stimulation on iridium oxide electrodes is highly variable between arrays, where one imaged array demonstrated noticeable stimulation-related damage and the other array had no such damage [29]. Furthermore, the study found that array recording quality was not compromised by stimulation. Similarly, other work did not show significant differences in SEM-visible degradation between platinum electrodes used for stimulation and iridium oxide electrodes used for stimulation [25, 26]. All of these previous studies used different stimulation protocols: a pulse train delivered at 20-300 Hz at amplitudes 1-100 *µ*A [29], a pulse train delivered at 300 Hz at amplitudes 1-210 *µ*A [25], and an undisclosed low-current stimulation [26]. Additionally, dissolution of electrodes due to current is most salient at aggressive levels of stimulation; one study reviewing electrode dissolution delivered variable amplitudes of current at 50 Hz for seven hours[42]; another study utilized currents at least one order of magnitude greater than the above protocols [39]. The electrolytic lesioning in this work primarily used direct currents delivered at around 150 *µ*A for 45 seconds, though parameters ranged between 50-450 *µ*A and 12-600 seconds. Thus, despite the differences between specific lesioning and stimulation protocols, the findings presented here are consistent with previous studies that found no unusual damage to the electrodes used for stimulation. This aligns with previous findings that electrolytic lesioning does not significantly impact the recording ability of electrodes [20].

The patterns of damage found in this work also largely reflect previous SEM analyses of explanted arrays. A prior study also found that the electrodes closest to the perimeter of the array suffered the most damage [26]. This pattern is also visually observable in other published results [27, 29]. This is likely because the edges of the arrays serve as a physical shield for the innermost electrodes. Edge electrodes are subject to greater access to brain tissue, as well as to mechanical damage incurred during surgery and handling.

The frequency of damage types identified in this work differ from previous studies, but is not unusual. In this work, the most common damage type within arrays used for lesioning is abnormal debris (n=276, 71.9%), followed by metal coating cracks (n=214, 55.7%), tip breakage (n=102, 26.6%), parylene C cracks (n=95, 24.7%), and parylene C delamination (n=48, 12.5%). On these lesioning arrays, there were n=53, or 13.8%, shank fractures. Similarly, across all eight NHP arrays imaged, the most common damage type is abnormal debris (n=518, 67.4%), followed by metal coating cracks (n=480, 62.5%), tip breakage (n=233, 30.3%), parylene C cracks (n=224, 29.2%), and parylene C delamination (n=60, 7.8%). Across all eight NHP arrays, there were n=117, or 15.2%, shank fractures. However, a prior study found that coating cracks were the most common type of observed damage, followed closely by parylene C cracks, then tip breakage, abnormal debris, and parylene C delamination [27]. While the frequency of damage types are not exactly aligned, previous work included a cleaning step during array preparation for SEM imaging; this work does not explicitly attempt to remove excess brain tissue from imaged arrays, which likely led to the relative increase we observed in abnormal debris. Additionally, variation in individual researcher’s scoring standards may also contribute to differences. As mentioned above, the overall spatial pattern of damage between these two studies is consistent. Finally, this study also found a moderate positive correlation between coating cracks and tip breakage. This reflects the fact that damage to the silicon core of an electrode often occurs in tandem with cracking and flaking of the surrounding metal coating; the silicon cannot break without also breaking its coating. This correlation is visible in the collected images presented in Figure 1.

## 5 Conclusion

This work analyzed high-definition SEM images of previously implanted electrode arrays used for electrolytic lesioning. Across all eight arrays previously implanted in NHPs, damage disproportionately occurred to the outer edges of arrays. Additionally, no statistically significant difference was found between the damage experienced by normal and lesioning electrodes within the same array. Furthermore, statistical testing between arrays used and not used for lesioning experiments did not indicate consistent, significant differences. Finally, this work also presents the largest publicly available set of SEM images of explanted arrays, consisting of eleven different multielectrode Utah arrays (ten 96-channel, one 64-channel) and one broken 96-channel array. These results from this dataset align with previous SEM studies of chronically implanted arrays used for both recording and stimulation, while providing further evidence for electrolytic lesioning as a safe and useful technique for experimental neuroscience.

## 6 Author Contributions

A Tor was responsible for data curation, formal analysis, investigation, methodology, software, validation, visualization, writ-ing, and review/editing. SE Clarke was responsible for conceptualization, data curation, supervision, validation, writing, and review/editing. IE Bray was responsible for conceptualization and review/editing. P Nuyujukian was responsible for con-ceptualization, funding acquisition, methodology, project administration, resources, supervision, writing, and review/editing. The members of the Brain Interfacing Laboratory provided additional support in review/editing.

## Acknowledgments

We thank S Baker for veterinary support, and K Chin and M Truong for administrative support. The members of the Brain Interfacing Laboratory who supported this work were Michelle S Wechsler, Mackenzie Risch, Alexandra Paraskevopoulou, Stephen I Ryu, Alissa S Ling, Elizabeth Jun, Michael P Silvernagel, Yuxin Wu, Kenji Y Marshall, Muhammad Abdulla, and Sydney Hunt. MS Wechsler, A Paraskevopoulou, K Lebedev, and MJ Risch were responsible for animal care and surgical support. SI Ryu was responsible for nonhuman primate array implantation. AS Ling, E Jun, MP Silvernagel, Y Wu, K Marshall, MU Abdulla, and S Hunt assisted in animal care. Part of this work was performed at the Stanford Nano Shared Facilities (SNSF), supported by the National Science Foundation under award ECCS-2026822. A Tor was supported by the Department of Defense (DoD) through the National Defense Science & Engineering Graduate (NDSEG) Fellowship Program and by a training grant from the National Science Foundation (1828993). SE Clarke was supported by a Stanford School of Medicine’s Dean’s Postdoctoral Fellowship. IE Bray was supported by an American Heart Association Predoctoral Fellowship (828653) and the National Science Foundation GRFP (1656518). This work was supported by a Stanford Human-Centered AI Seed Research Grant to SE Clarke and P Nuyujukian. This work was additionally supported by the following to P Nuyujukian: the National Institutes of Health (R01NS123517, R01NS130789, U19NS118284) and the Stanford Wu Tsai Neurosciences Institute.

## Supplemental Materials

**Supplemental Figure 1:**
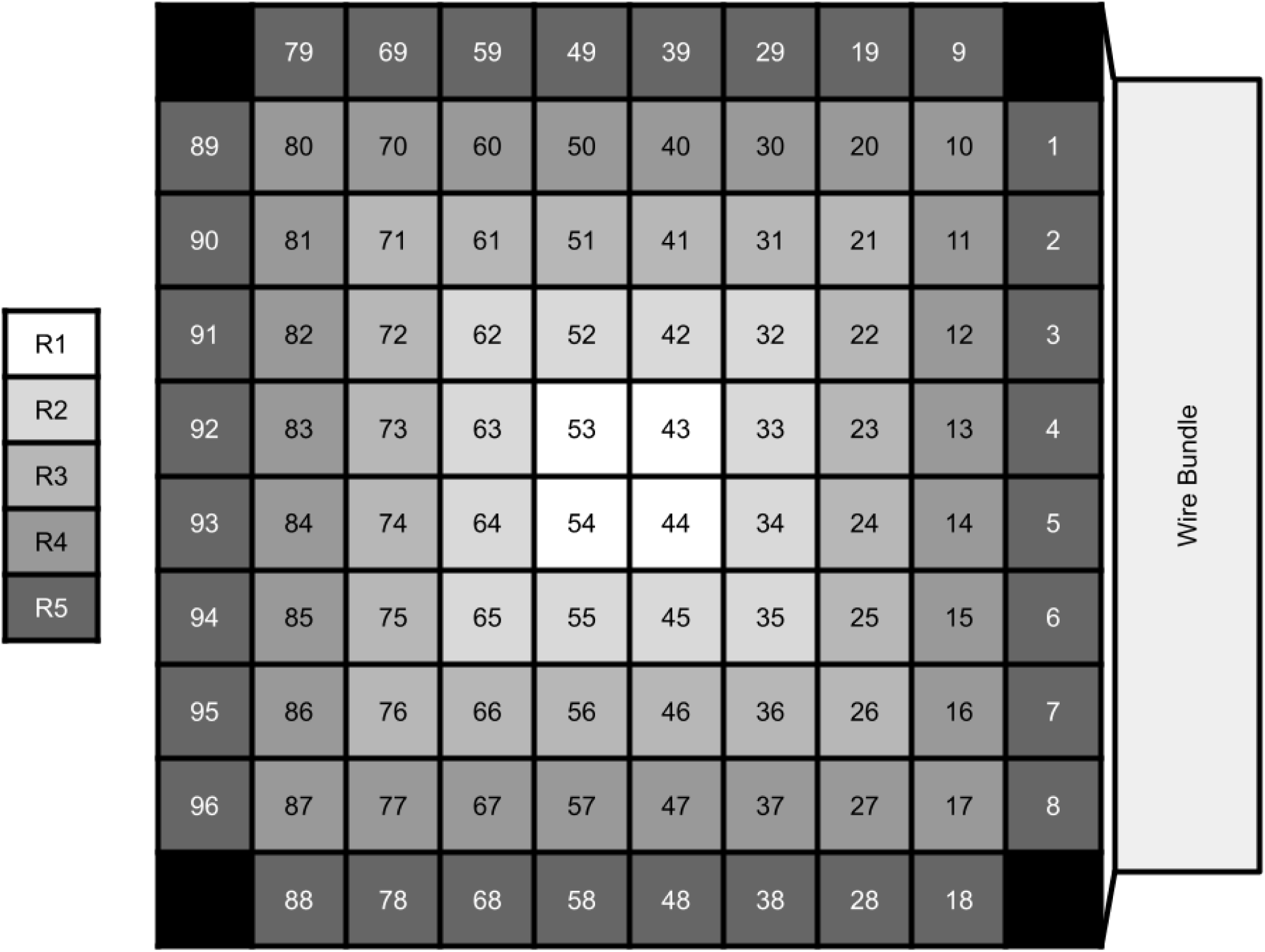
Numbering layout of the electrodes as imaged (pins facing outwards, towards the reader). Wire bundle is arranged to the right. Each radial section of the array is also color-coded and labeled. R1 refers to the innermost core electrodes of the array, R2 refers to the next outer ring of the array, and so on.

**Supplemental Figure 2:**
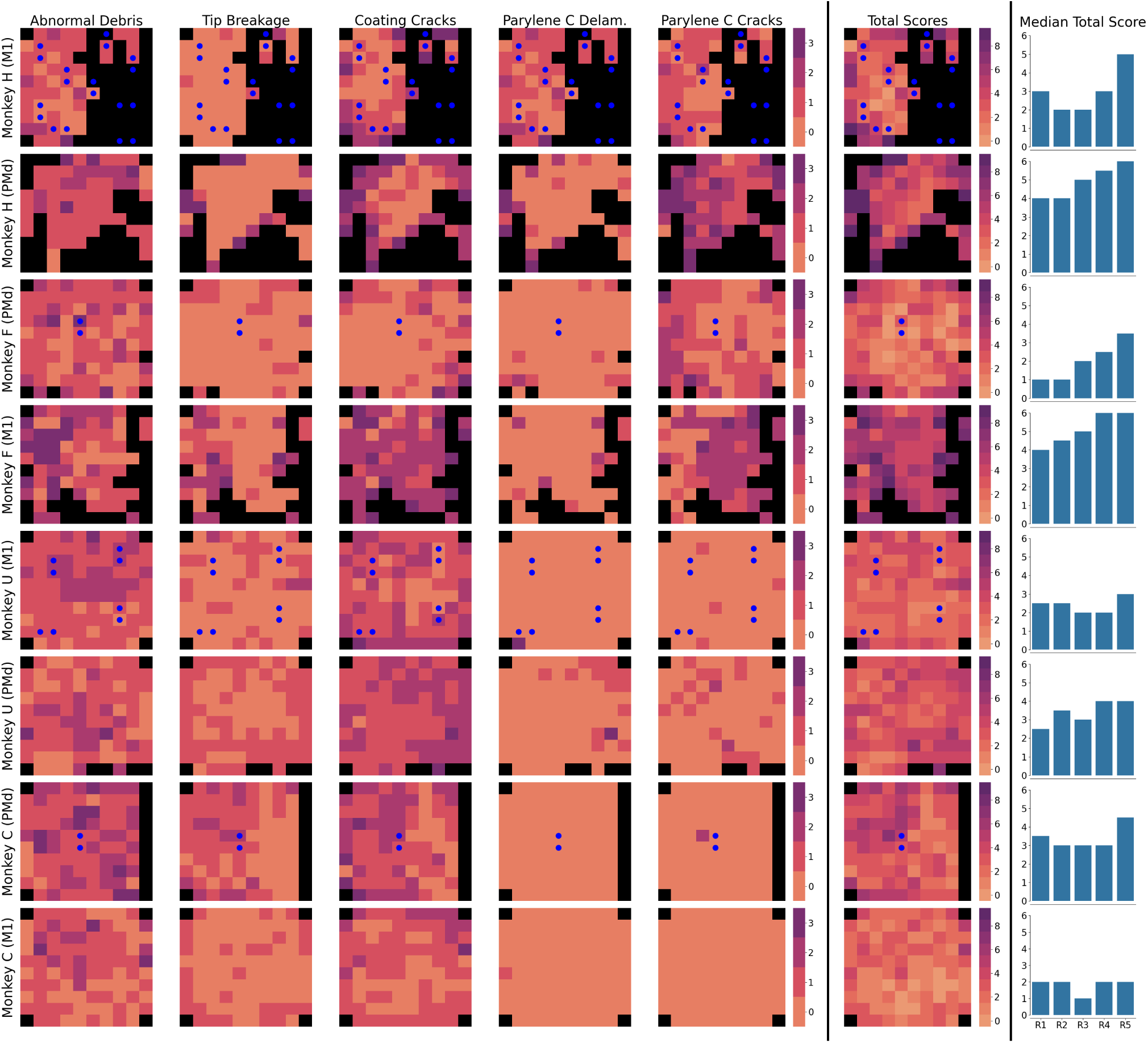
Heatmaps of damage scores (0-3) across the five identified types of damage and across all eight intact, imaged NHP arrays. Electrodes are displayed using the orientation in Figure 1 (electrode tips facing viewer, wire bundle on the right). Second-to-rightmost column displays summed damage scores for each array across the five types of damage. Electrodes used for electrolytic lesioning are denoted with blue dots. Median summed scores for each radial section of the array are plotted in the bar charts to the right of the heatmaps. Ring layout and numbering information is available in Supplementary Figure 1. Unwired electrodes (electrodes not wire bonded at time of manufacture) and electrodes with shank fractures are ignored and displayed in black, as they are not scored.

**Supplemental Figure 3:**
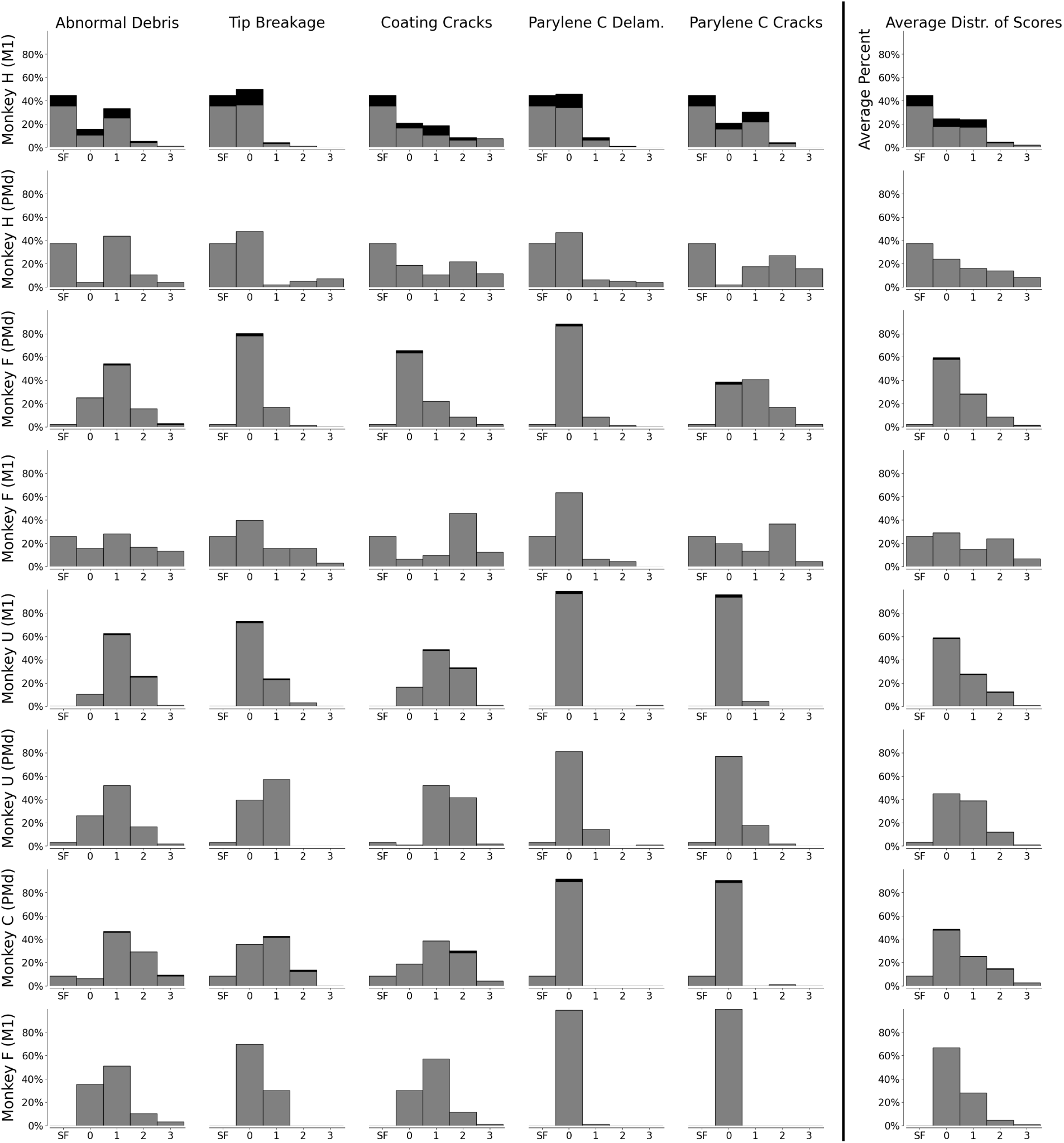
Stacked histograms of damage scores (0-3) across the five identified types of damage and across all eight intact, NHP imaged arrays. Gray indicates normal electrodes, and black indicates lesioning electrodes. Rightmost column displays the average distribution of damage scores across the five types of damage. Electrodes with shank fractures (SF) are ignored, as they are not scored.

**Supplemental Figure 4:**
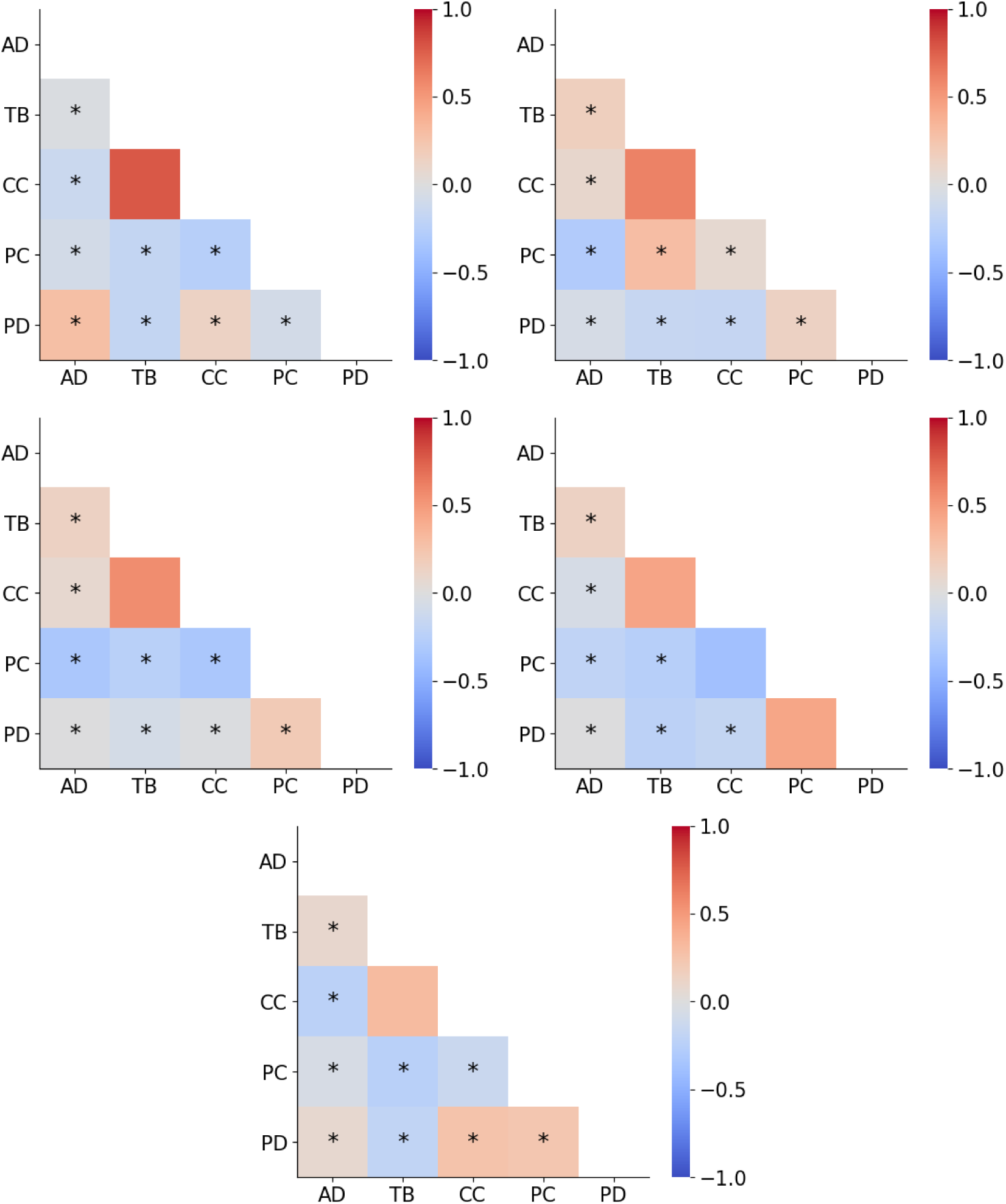
Correlation plots (Pearson’s R) for each of the five rings across all four imaged lesioning arrays. Test values with Bonferroni-corrected *p <* 0.05 are displayed with asterisks. Raw *r* and *p*-values are separately available in Supplemental Table 5.

**Supplemental Figure 5:**
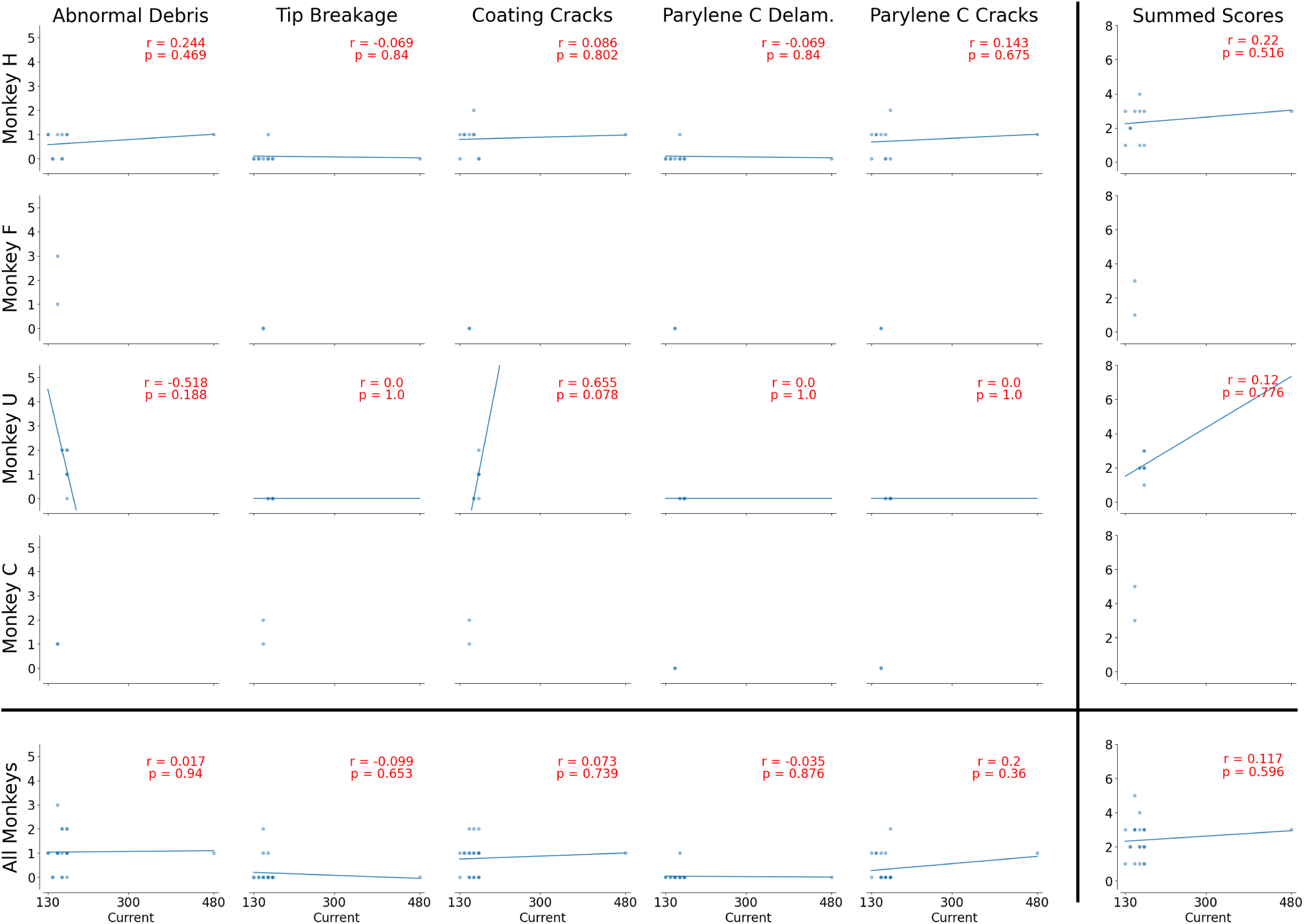
Scatterplots of the lesion electrodes’ damage scores over the target current for each lesion. A best-fit line is also plotted along with regression results. In cases where only one target current was used (ie. subjects with only one lesion), linear regression was not performed. No best-fit lines were statistically significant at *p* = 0.05.

**Supplemental Figure 6:**
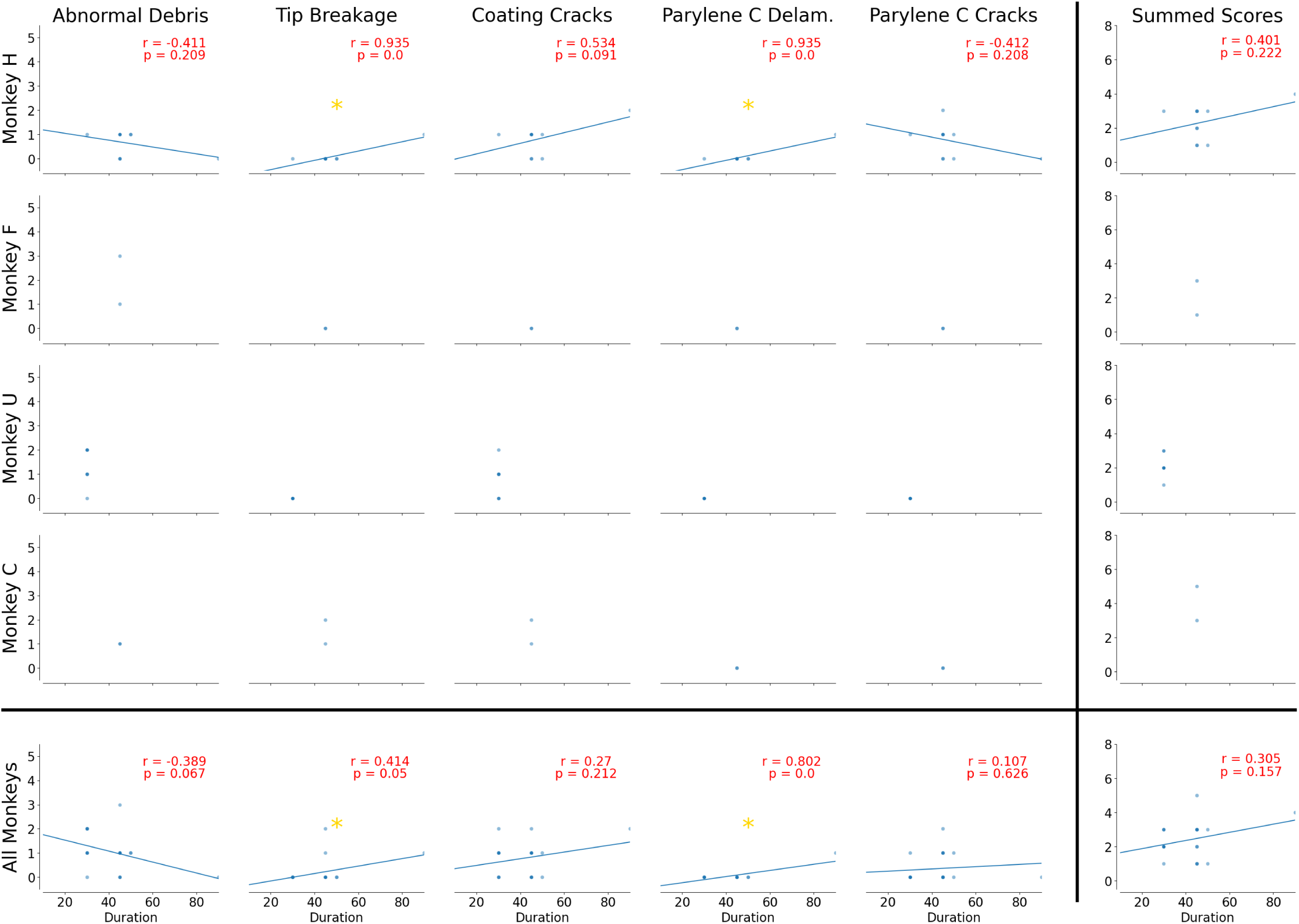
Scatterplots of the lesion electrodes’ damage scores over the target lesioning procedure duration for each lesion. A best-fit line is also plotted along with regression results. In cases where only one duration was used (ie. subjects with only one lesion, subjects with the same duration across all lesions), linear regression was not performed. Scatterplots with statistically significant lines of best fit are starred in yellow.

**Supplemental Figure 7:**
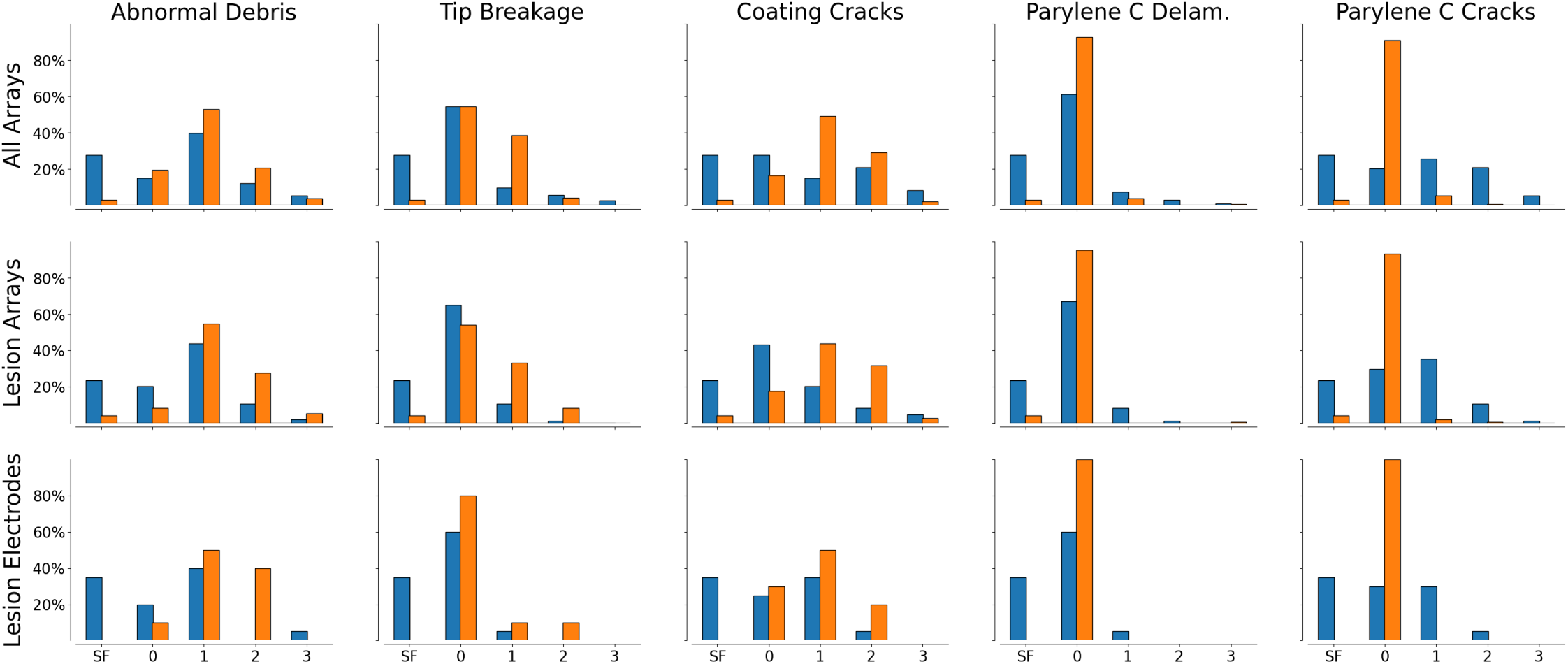
Histograms of the lesion electrodes’ damage scores, separated by array material. Blue bars are IrOx arrays and orange bars are Pt arrays. The first row shows scores across all eight scored NHP arrays (n=384 IrOx electrodes, n=384 Pt electrodes). The second row shows scores for only the four NHP arrays used for lesioning (n=192 IrOx electrodes, n=192 Pt electrodes). The third row shows scores for only the subset of electrodes used for NHP lesioning (n=10 IrOx electrodes, n=20 Pt electrodes). Notably, it appears that platinum arrays imaged in this work generally appear to report less damage than iridium oxide arrays when considering all arrays. Additionally, the imaged platinum arrays appear to always report less Parylene C damage. However, platinum arrays generally appear to report more other types of damage (abnormal debris, tip breakage, coating cracks) when used for lesioning. This supports previous literature that indicates IrOx may be more resistant to stimulation-related damage than Pt [26, 41–43]. It is important to note that many other factors, such as time in tissue, explant surgery, and immune response of the specific implant subject, all impact the conclusions of this analysis.

**Supplemental Table 1:** Details and characteristics for all imaged and analyzed devices in this work.

| Subject | Region | Metal | Implant Date | Explant Date | Days | Serial Num | Size | Notes |
| --- | --- | --- | --- | --- | --- | --- | --- | --- |
| H | PMd | Pt | Mar 17 2014 | 6 May 2020 | 2242 | 1024-1177 | 96 | Missing wire bundle – layout/numbering is arbitrary |
| H | M1 | Pt | Mar 17 2014 | 6 May 2020 | 2242 | 1024-1166 | 96 |  |
| F | PMd | Pt | Sep 8 2014 | 27 Oct 2019 | 1875 | 1024-1171 | 96 |  |
| F | M1 | Pt | Sep 8 2014 | 27 Oct 2019 | 1875 | 1024-1175 | 96 | Missing wire bundle; previously encapsulated in fibrin |
| U | PMd | IrOx | Aug 4 2017 | Dec 5 2024 | 2680 | 1024-1903 | 96 |  |
| U | M1 medial posterior | IrOx | Aug 4 2017 | Dec 5 2024 | 2680 | 1024-1902 | 96 | Broken during extraction |
| U | M1 lateral anterior | IrOx | Aug 4 2017 | Dec 5 2024 | 2680 | 1024-1905 | 96 |  |
| C | PMd | IrOx | Mar 25 2021 | 9 Nov 2022 | 594 | 6250-001604 | 96 |  |
| C | M1 | IrOx | Mar 25 2021 | 9 Nov 2022 | 594 | 6250-001608 | 96 |  |
| Agar | x |  | 25 Aug 2022 | 25 Aug 2022 | x | 1025-149 | 64 | Implanted briefly into agar gel and used to test initial electrolytic lesioning protocols during development. |
| P | x | Pt | x | x | x | 1024-0577 | 96 | Used in multiple lesions associated with Bray*, Clarke*, et al., eLife 2024. Implanted and removed from brain tissue multiple times over weeks/months. |
| Control | x | x | x | x | x |  | 96 | Never-implanted array. Damage and debris are due to handling. One image is available per column. |

**Supplemental Table 2:** Shapiro-Wilkes p-values for different electrode populations.

| Subject | Region | Lesioning Array | Population Type | Damage Type | Shapiro-Wilk p-value |
| --- | --- | --- | --- | --- | --- |
| H | M1 | x | B | AD | < 0.001 |
| H | M1 | x | L | AD | < 0.001 |
| H | M1 | x | N | AD | < 0.001 |
| H | M1 | x | B | TB | < 0.001 |
| H | M1 | x | L | TB | < 0.001 |
| H | M1 | x | N | TB | < 0.001 |
| H | M1 | x | B | CC | < 0.001 |
| H | M1 | x | L | CC | < 0.01 |
| H | M1 | x | N | CC | < 0.001 |
| H | M1 | x | B | PC | < 0.001 |
| H | M1 | x | L | PC | < 0.01 |
| H | M1 | x | N | PC | < 0.001 |
| H | M1 | x | B | PD | < 0.001 |
| H | M1 | x | L | PD | < 0.001 |
| H | M1 | x | N | PD | < 0.001 |
| H | M1 | x | B | TO | < 0.01 |
| H | M1 | x | L | TO | 5.439E-002 |
| H | M1 | x | N | TO | < 0.01 |
| H | PMd |  | N | AD | < 0.001 |
| H | PMd |  | N | TB | < 0.001 |
| H | PMd |  | N | CC | < 0.001 |
| H | PMd |  | N | PC | < 0.001 |
| H | PMd |  | N | PD | < 0.001 |
| H | PMd |  | N | TO | < 0.001 |
| F | PMd | x | B | AD | < 0.001 |
| F | PMd | x | N | AD | < 0.001 |
| F | PMd | x | B | TB | < 0.001 |
| F | PMd | x | N | TB | < 0.001 |
| F | PMd | x | B | CC | < 0.001 |
| F | PMd | x | N | CC | < 0.001 |
| F | PMd | x | B | PC | < 0.001 |
| F | PMd | x | N | PC | < 0.001 |
| F | PMd | x | B | PD | < 0.001 |
| F | PMd | x | N | PD | < 0.001 |
| F | PMd | x | B | TO | < 0.001 |
| F | PMd | x | N | TO | < 0.01 |
| F | M1 |  | N | AD | < 0.001 |
| F | M1 |  | N | TB | < 0.001 |
| F | M1 |  | N | CC | < 0.001 |
| F | M1 |  | N | PC | < 0.001 |
| F | M1 |  | N | PD | < 0.001 |
| F | M1 |  | N | TO | < 0.01 |
| U | M1 | x | B | AD | < 0.001 |
| U | M1 | x | L | AD | < 0.05 |
| U | M1 | x | N | AD | < 0.001 |
| U | M1 | x | B | TB | < 0.001 |
| U | M1 | x | L | TB | 1.000E+000 |
| U | M1 | x | N | TB | < 0.001 |
| U | M1 | x | B | CC | < 0.001 |
| U | M1 | x | L | CC | 5.552E-002 |
| U | M1 | x | N | CC | < 0.001 |
| U | M1 | x | B | PC | < 0.001 |
| U | M1 | x | L | PC | 1.000E+000 |
| U | M1 | x | N | PC | < 0.001 |
| U | M1 | x | B | PD | < 0.001 |
| U | M1 | x | L | PD | 1.000E+000 |
| U | M1 | x | N | PD | < 0.001 |
| U | M1 | x | B | TO | < 0.001 |
| U | M1 | x | L | TO | < 0.05 |
| U | M1 | x | N | TO | < 0.001 |
| U | PMd |  | N | AD | < 0.001 |
| U | PMd |  | N | TB | < 0.001 |
| U | PMd |  | N | CC | < 0.001 |
| U | PMd |  | N | PC | < 0.001 |
| U | PMd |  | N | PD | < 0.001 |
| U | PMd |  | N | TO | < 0.001 |
| C | PMd | x | B | AD | < 0.001 |
| C | PMd | x | N | AD | < 0.001 |
| C | PMd | x | B | TB | < 0.001 |
| C | PMd | x | N | TB | < 0.001 |
| C | PMd | x | B | CC | < 0.001 |
| C | PMd | x | N | CC | < 0.001 |
| C | PMd | x | B | PC | < 0.001 |
| C | PMd | x | N | PC | < 0.001 |
| C | PMd | x | B | PD | 1.000E+000 |
| C | PMd | x | N | PD | 1.000E+000 |
| C | PMd | x | B | TO | < 0.001 |
| C | PMd | x | N | TO | < 0.001 |
| C | M1 |  | N | AD | < 0.001 |
| C | M1 |  | N | TB | < 0.001 |
| C | M1 |  | N | CC | < 0.001 |
| C | M1 |  | N | PC | 1.000E+000 |
| C | M1 |  | N | PD | < 0.001 |
| C | M1 |  | N | TO | < 0.001 |

**Supplemental Table 3:** Mann-Whitney U and Levene test p-values when comparing scores between lesioning electrodes and non-lesioning elec-trodes on the same array.

| Subject | Damage Type | Mann-Whitney p-value | Levene p-value |
| --- | --- | --- | --- |
| C | AD | 3.49E-01 | 2.10E-01 |
| C | TB | 1.46E-01 | 9.22E-01 |
| C | CC | 6.02E-01 | 7.55E-01 |
| C | PC | 9.39E-01 | 8.80E-01 |
| C | PD | 1.00E+00 |  |
| C | TO | 5.87E-01 | 7.71E-01 |
| H | AD | 2.67E-01 | 7.24E-01 |
| H | TB | 9.66E-01 | 8.27E-01 |
| H | CC | 7.12E-01 | < 0.05 |
| H | PC | 8.82E-01 | 9.90E-01 |
| H | PD | 4.39E-01 | 4.14E-01 |
| H | TO | 5.77E-01 | 9.19E-02 |
| F | AD | 1.62E-01 | 1.72E-01 |
| F | TB | 5.17E-01 | 5.19E-01 |
| F | CC | 3.35E-01 | 3.82E-01 |
| F | PC | 1.11E-01 | 1.20E-01 |
| F | PD | 6.62E-01 | 6.60E-01 |
| F | TO | 6.32E-01 | 6.50E-01 |
| U | AD | 2.76E-01 | 1.58E-01 |
| U | TB | 7.62E-02 | 8.96E-02 |
| U | CC | 7.66E-02 | 9.07E-01 |
| U | PC | 5.53E-01 | 5.43E-01 |
| U | PD | 7.92E-01 | 7.65E-01 |
| U | TO | 6.59E-02 | 1.08E-01 |
| Aggregate | AD | 4.81E-01 | 9.52E-01 |
| Aggregate | TB | 6.82E-02 | 1.08E-01 |
| Aggregate | CC | 3.89E-01 | 8.39E-02 |
| Aggregate | PC | 7.62E-01 | 6.76E-01 |
| Aggregate | PD | 9.28E-01 | 9.08E-01 |
| Aggregate | TO | 1.05E-01 | 6.31E-02 |

**Supplemental Table 4:**
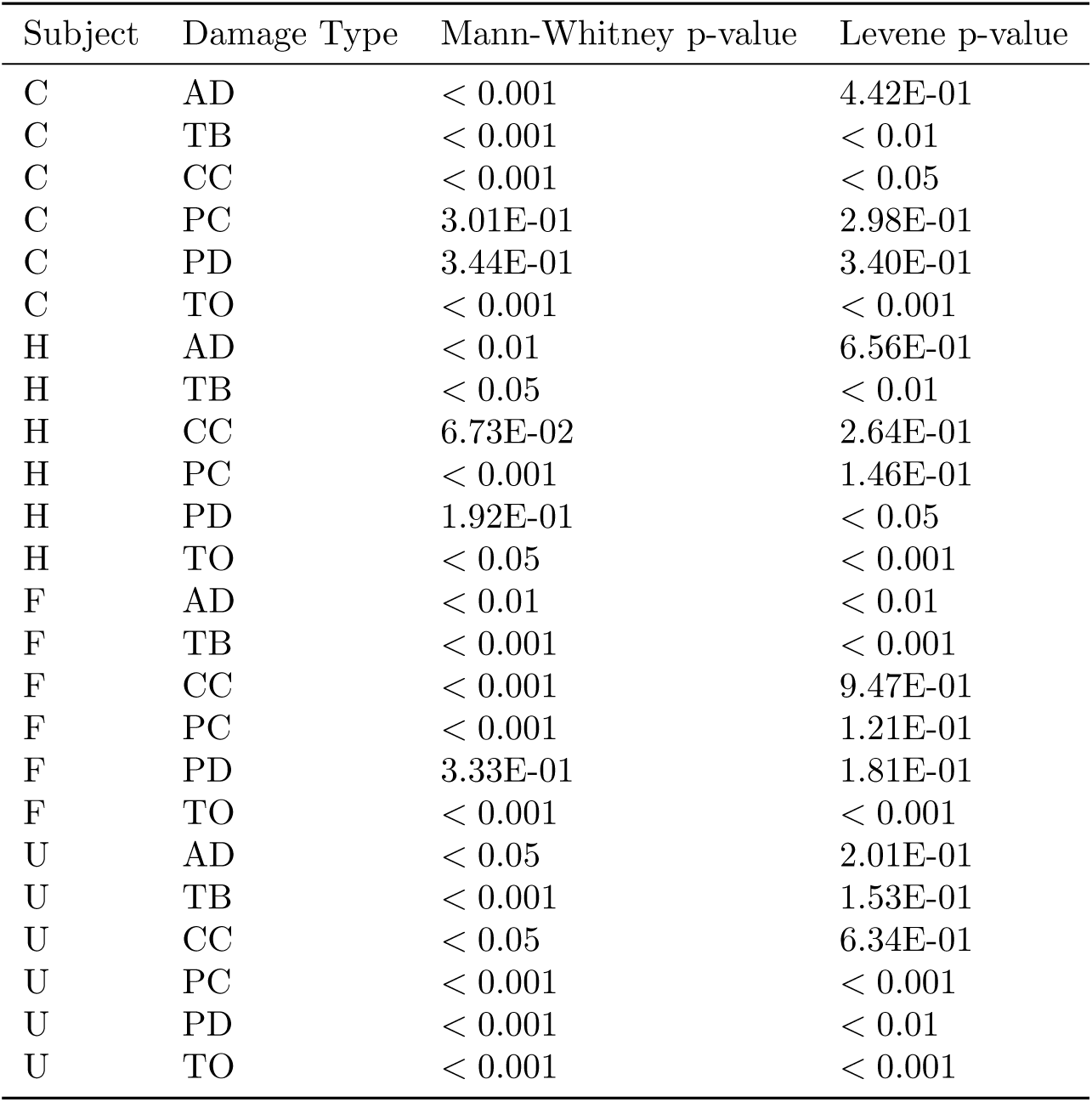
Mann-Whitney U and Levene test p-values when comparing scores between all electrodes on arrays used and not used for lesioning experiments when implanted in the same subject.

**Supplemental Table 5:** Bonferroni-corrected p-value is 0.005 (0.05/n=10 comparisons).

|  | All |  | R1 |  | R2 |  | R3 |  | R4 |  | R5 |  |
| --- | --- | --- | --- | --- | --- | --- | --- | --- | --- | --- | --- | --- |
| Damage Types | r | p-val | r | p-val | r | p-val | r | p-val | r | p-val | r | p-val |
| AD/AD | 1.0000 | < 0.001 | 1.0000 | < 0.001 | 1.0000 | < 0.001 | 1.0000 | < 0.001 | 1.0000 | < 0.001 | 1.0000 | < 0.001 |
| AD/TB | 0.1226 | < 0.05 | -0.0292 | 0.9178 | 0.1591 | 0.3141 | 0.1387 | 0.2521 | 0.1375 | 0.1704 | 0.0792 | 0.4268 |
| AD/CC | -0.0443 | 0.4215 | -0.1274 | 0.6509 | 0.0937 | 0.5549 | 0.0778 | 0.5221 | -0.0619 | 0.5383 | -0.2294 | 0.0198 |
| AD/PC | -0.1621 | < 0.005 | -0.0714 | 0.8003 | -0.2846 | 0.0677 | -0.3252 | < 0.01 | -0.1970 | < 0.05 | -0.0499 | 0.6163 |
| AD/PD | 0.0437 | 0.4278 | 0.2857 | 0.3019 | -0.0577 | 0.7168 | -0.0018 | 0.9882 | -0.0064 | 0.9492 | 0.0775 | 0.4366 |
| TB/TB | 1.0000 | < 0.001 | 1.0000 | < 0.001 | 1.0000 | < 0.001 | 1.0000 | < 0.001 | 1.0000 | < 0.001 | 1.0000 | < 0.001 |
| TB/CC | 0.4676 | < 0.001 | 0.7802 | < 0.001 | 0.6069 | < 0.001 | 0.5575 | < 0.001 | 0.4405 | < 0.001 | 0.3100 | < 0.005 |
| TB/PC | -0.1527 | < 0.01 | -0.1750 | 0.5328 | 0.2901 | 0.0624 | -0.2391 | < 0.05 | -0.2535 | < 0.05 | -0.2411 | < 0.05 |
| TB/PD | -0.1449 | < 0.01 | -0.1750 | 0.5328 | -0.1614 | 0.3071 | -0.0674 | 0.5792 | -0.2227 | < 0.05 | -0.1925 | 0.0514 |
| CC/CC | 1.0000 | < 0.001 | 1.0000 | < 0.001 | 1.0000 | < 0.001 | 1.0000 | < 0.001 | 1.0000 | < 0.001 | 1.0000 | < 0.001 |
| CC/PC | -0.1366 | < 0.05 | -0.2548 | 0.3594 | 0.0677 | 0.6701 | -0.3252 | < 0.01 | -0.3775 | < 0.001 | -0.1396 | 0.1595 |
| CC/PD | 0.1203 | < 0.05 | 0.1274 | 0.6509 | -0.1658 | 0.2940 | -0.0188 | 0.8775 | -0.1760 | 0.0783 | 0.2562 | < 0.01 |
| PC/PC | 1.0000 | < 0.001 | 1.0000 | < 0.001 | 1.0000 | < 0.001 | 1.0000 | < 0.001 | 1.0000 | < 0.001 | 1.0000 | < 0.001 |
| PC/PD | 0.2889 | < 0.001 | -0.0714 | 0.8003 | 0.1332 | 0.4003 | 0.1979 | 0.1005 | 0.4326 | < 0.001 | 0.2292 | < 0.05 |
| PD/PD | 1.0000 | < 0.001 | 1.0000 | < 0.001 | 1.0000 | < 0.001 | 1.0000 | < 0.001 | 1.0000 | < 0.001 | 1.0000 | < 0.001 |

**Supplemental Table 6:**
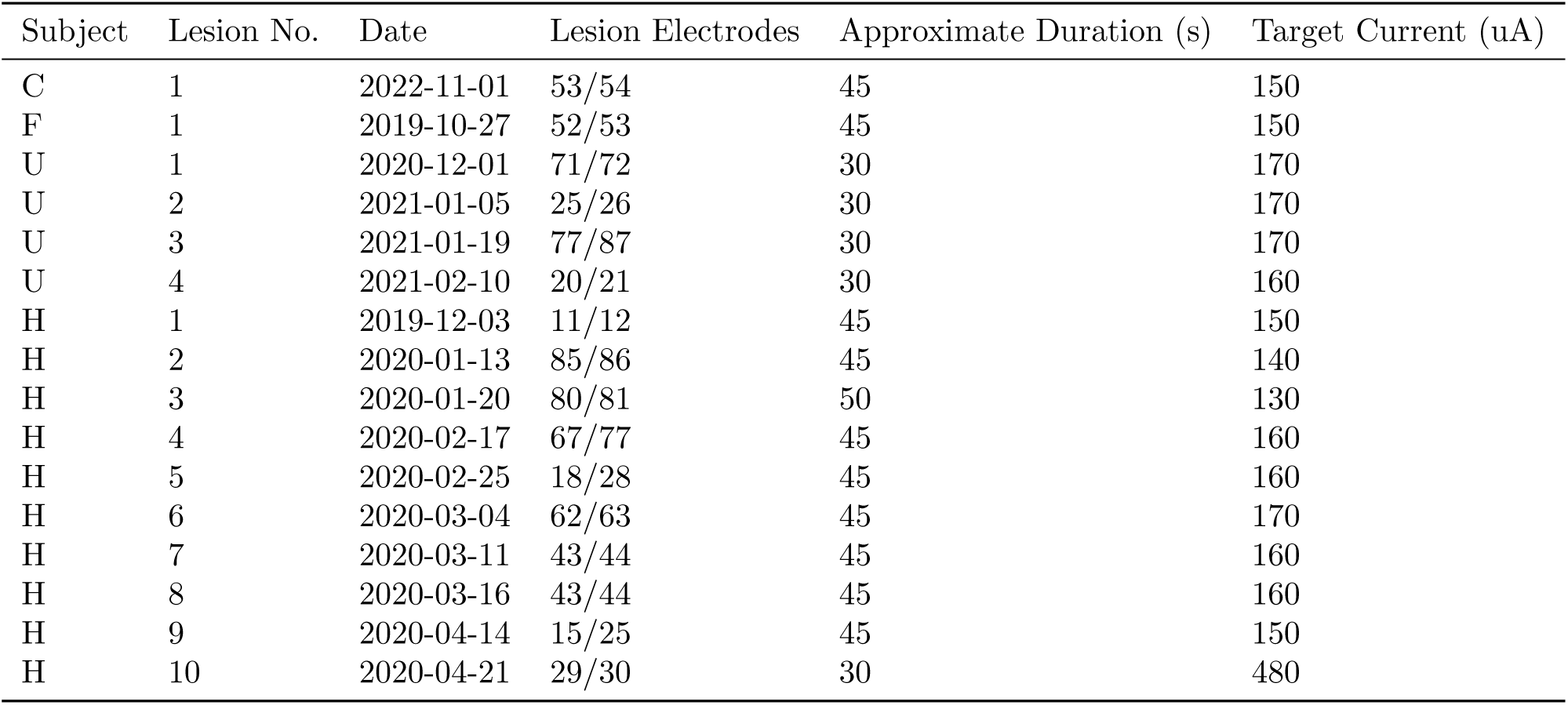
Lesion parameters for NHP lesions analyzed in this work.

## References

[1] J. P. Cunningham and B. M. Yu, “Dimensionality reduction for large-scale neural recordings,” Nature Neuroscience, vol. 17, p. 1500–1509, Aug. 2014.

[2] S. Vyas, M. D. Golub, D. Sussillo, and K. V. Shenoy, “Computation through neural population dynamics,” Annual Review of Neuroscience, vol. 43, p. 249–275, July 2020.

[3] R. B. Ebitz and B. Y. Hayden, “The population doctrine in cognitive neuroscience,” Neuron, vol. 109, p. 3055–3068, Oct. 2021.

[4] E. M. Maynard, C. T. Nordhausen, and R. A. Normann, “The utah intracortical electrode array: A recording structure for potential brain-computer interfaces,” Electroencephalography and Clinical Neurophysiology, vol. 102, p. 228–239, Mar. 1997.

[5] M. D. Serruya, N. G. Hatsopoulos, L. Paninski, M. R. Fellows, and J. P. Donoghue, “Instant neural control of a movement signal,” Nature, vol. 416, p. 141–142, Mar. 2002.

[6] V. Gilja, P. Nuyujukian, C. A. Chestek, J. P. Cunningham, B. M. Yu, J. M. Fan, M. M. Churchland, M. T. Kaufman, J. C. Kao, S. I. Ryu, and K. V. Shenoy, “A high-performance neural prosthesis enabled by control algorithm design,” Nature Neuroscience, vol. 15, p. 1752–1757, Nov. 2012.

[7] M. M. Churchland, J. P. Cunningham, M. T. Kaufman, J. D. Foster, P. Nuyujukian, S. I. Ryu, and K. V. Shenoy, “Neural population dynamics during reaching,” Nature, vol. 487, p. 51–56, June 2012.

[8] J. A. Gallego, M. G. Perich, R. H. Chowdhury, S. A. Solla, and L. E. Miller, “Long-term stability of cortical population dynamics underlying consistent behavior,” Nature Neuroscience, vol. 23, p. 260–270, Jan. 2020.

[9] V. Gilja, C. Pandarinath, C. H. Blabe, P. Nuyujukian, J. D. Simeral, A. A. Sarma, B. L. Sorice, J. A. Perge, B. Jarosiewicz, L. R. Hochberg, K. V. Shenoy, and J. M. Henderson, “Clinical translation of a high-performance neural prosthesis,” Nature Medicine, vol. 21, p. 1142–1145, Sept. 2015.

[10] F. R. Willett, E. M. Kunz, C. Fan, D. T. Avansino, G. H. Wilson, E. Y. Choi, F. Kamdar, M. F. Glasser, L. R. Hochberg, S. Druckmann, K. V. Shenoy, and J. M. Henderson, “A high-performance speech neuroprosthesis,” Nature, vol. 620, p. 1031–1036, Aug. 2023.

[11] P. R. Roelfsema and S. Treue, “Basic neuroscience research with nonhuman primates: A small but indispensable component of biomedical research,” Neuron, vol. 82, p. 1200–1204, June 2014.

[12] A. S. Mitchell, A. Thiele, C. I. Petkov, A. Roberts, T. W. Robbins, W. Schultz, and R. Lemon, “Continued need for non-human primate neuroscience research,” Current Biology, vol. 28, p. R1186–R1187, Oct. 2018.

[13] E. J. Tehovnik, A. S. Tolias, F. Sultan, W. M. Slocum, and N. K. Logothetis, “Direct and indirect activation of cortical neurons by electrical microstimulation,” Journal of Neurophysiology, vol. 96, p. 512–521, Aug. 2006.

[14] M. T. Kline, “chapter 169 - radiofrequency techniques,” in *Pain Management* (S. D. Waldman and J. I. Bloch, eds.), pp. 1411–1459, Philadelphia: W.B. Saunders, 2007.

[15] V. HORSLEY and R. H. Clarke, “The structure and functions of the cerebellum examined by a new method.,” Brain, vol. 31, no. 1, p. 45–124, 1908.

[16] W. H. SWEET, “Unipolar anodal electrolytic lesions in the brain of man and cat: Report of five human cases with electrically produced bulbar or mesencephalic tractotomies,” *A.M.A. Archives of Neurology & Psychiatry*, vol. 70, p. 224, Aug. 1953.

[17] Y.-Y. Chen, H.-Y. Lai, S.-H. Lin, C.-W. Cho, W.-H. Chao, C.-H. Liao, S. Tsang, Y.-F. Chen, and S.-Y. Lin, “Design and fabrication of a polyimide-based microelectrode array: Application in neural recording and repeatable electrolytic lesion in rat brain,” Journal of Neuroscience Methods, vol. 182, p. 6–16, Aug. 2009.

[18] G. Townsend, P. Peloquin, F. Kloosterman, J. F. Hetke, and L. Leung, “Recording and marking with silicon multichannel electrodes,” Brain Research Protocols, vol. 9, p. 122–129, Apr. 2002.

[19] B. Chehrazi and W. F. Collins, “A comparison of effects of bipolar and monopolar electrocoagulation in brain,” Journal of Neurosurgery, vol. 54, p. 197–203, Feb. 1981.

[20] I. E. Bray, S. E. Clarke, K. M. Casey, and P. Nuyujukian, “Neuroelectrophysiology-compatible electrolytic lesioning,” eLife, vol. 12, July 2024.

[21] S. Suner, M. Fellows, C. Vargas-Irwin, G. Nakata, and J. Donoghue, “Reliability of signals from a chronically implanted, silicon-based electrode array in non-human primate primary motor cortex,” IEEE Transactions on Neural Systems and Rehabilitation Engineering, vol. 13, p. 524–541, Dec. 2005.

[22] J. D. Simeral, S.-P. Kim, M. J. Black, J. P. Donoghue, and L. R. Hochberg, “Neural control of cursor trajectory and click by a human with tetraplegia 1000 days after implant of an intracortical microelectrode array,” Journal of Neural Engineering, vol. 8, p. 025027, Mar. 2011.

[23] J. E. Downey, N. Schwed, S. M. Chase, A. B. Schwartz, and J. L. Collinger, “Intracortical recording stability in human brain–computer interface users,” Journal of Neural Engineering, vol. 15, p. 046016, May 2018.

[24] C. A. Chestek, V. Gilja, P. Nuyujukian, J. D. Foster, J. M. Fan, M. T. Kaufman, M. M. Churchland, Z. Rivera-Alvidrez, J. P. Cunningham, S. I. Ryu, and K. V. Shenoy, “Long-term stability of neural prosthetic control signals from silicon cortical arrays in rhesus macaque motor cortex,” Journal of Neural Engineering, vol. 8, p. 045005, July 2011.

[25] X. Chen, F. Wang, R. Kooijmans, P. C. Klink, C. Boehler, M. Asplund, and P. R. Roelfsema, “Chronic stability of a neuroprosthesis comprising multiple adjacent utah arrays in monkeys,” Journal of Neural Engineering, vol. 20, p. 036039, June 2023.

[26] D. A. Bjanes, S. Kellis, R. Nickl, B. Baker, T. Aflalo, L. Bashford, S. Chivukula, M. S. Fifer, L. E. Osborn, B. Christie, B. A. Wester, P. A. Celnik, D. Kramer, K. Pejsa, N. E. Crone, W. S. Anderson, N. Pouratian, B. Lee, C. Y. Liu, F. Tenore, L. Rieth, and R. A. Andersen, “Quantifying physical degradation alongside recording and stimulation performance of 980 intracortical microelectrodes chronically implanted in three humans for 956-2246 days,” Sept. 2024.

[27] P. R. Patel, E. J. Welle, J. G. Letner, H. Shen, A. J. Bullard, C. M. Caldwell, A. Vega-Medina, J. M. Richie, H. E. Thayer, P. G. Patil, D. Cai, and C. A. Chestek, “Utah array characterization and histological analysis of a multi-year implant in non-human primate motor and sensory cortices,” Journal of Neural Engineering, vol. 20, p. 014001, Jan. 2023.

[28] J. C. Barrese, J. Aceros, and J. P. Donoghue, “Scanning electron microscopy of chronically implanted intracortical microelectrode arrays in non-human primates,” Journal of Neural Engineering, vol. 13, p. 026003, Jan. 2016.

[29] K. Woeppel, C. Hughes, A. J. Herrera, J. R. Eles, E. C. Tyler-Kabara, R. A. Gaunt, J. L. Collinger, and X. T. Cui, “Explant analysis of utah electrode arrays implanted in human cortex for brain-computer-interfaces,” Frontiers in Bioengineering and Biotechnology, vol. 9, Dec. 2021.

[30] D. Yi, Y. Yao, Y. Wang, and L. Chen, “Manufacturing processes of implantable microelectrode array for in vivo neural electrophysiological recordings and stimulation: A state-of-the-art review,” Journal of Micro- and Nano-Manufacturing, vol. 10, Dec. 2022.

[31] P. Campbell, K. Jones, R. Huber, K. Horch, and R. Normann, “A silicon-based, three-dimensional neural interface: manufacturing processes for an intracortical electrode array,” IEEE Transactions on Biomedical Engineering, vol. 38, no. 8, p. 758–768, 1991.

[32] R. Bhandari, S. Negi, and F. Solzbacher, “Wafer-scale fabrication of penetrating neural microelectrode arrays,” Biomedical Microdevices, vol. 12, p. 797–807, May 2010.

[33] R. Caldwell, M. G. Street, R. Sharma, P. Takmakov, B. Baker, and L. Rieth, “Characterization of parylene-c degradation mechanisms: In vitro reactive accelerated aging model compared to multiyear in vivo implantation,” Biomaterials, vol. 232, p. 119731, Feb. 2020.

[34] J. C. Barrese, N. Rao, K. Paroo, C. Triebwasser, C. Vargas-Irwin, L. Franquemont, and J. P. Donoghue, “Failure mode analysis of silicon-based intracortical microelectrode arrays in non-human primates,” Journal of Neural Engineering, vol. 10, p. 066014, Nov. 2013.

[35] V. S. Polikov, P. A. Tresco, and W. M. Reichert, “Response of brain tissue to chronically implanted neural electrodes,” Journal of Neuroscience Methods, vol. 148, p. 1–18, Oct. 2005.

[36] J. W. Salatino, K. A. Ludwig, T. D. Y. Kozai, and E. K. Purcell, “Glial responses to implanted electrodes in the brain,” Nature Biomedical Engineering, vol. 1, p. 862–877, Nov. 2017.

[37] R. Biran, D. C. Martin, and P. A. Tresco, “The brain tissue response to implanted silicon microelectrode arrays is increased when the device is tethered to the skull,” Journal of Biomedical Materials Research Part A, vol. 82A, p. 169–178, May 2007.

[38] K. A. Potter, A. C. Buck, W. K. Self, and J. R. Capadona, “Stab injury and device implantation within the brain results in inversely multiphasic neuroinflammatory and neurodegenerative responses,” Journal of Neural Engineering, vol. 9, p. 046020, July 2012.

[39] R. K. Shepherd, P. M. Carter, A. N. Dalrymple, Y. L. Enke, A. K. Wise, T. Nguyen, J. Firth, A. Thompson, and J. B. Fallon, “Platinum dissolution and tissue response following long-term electrical stimulation at high charge densities,” Journal of Neural Engineering, vol. 18, p. 036021, Mar. 2021.

[40] D. D. Shah, P. Carter, M. N. Shivdasani, N. Fong, W. Duan, D. Esrafilzadeh, L. A. Poole-Warren, and U. A. Aregueta Robles, “Deciphering platinum dissolution in neural stimulation electrodes: Electrochemistry or biology?,” Biomaterials, vol. 309, p. 122575, Sept. 2024.

[41] S. Cogan, T. Plante, and J. Ehrlich, “Sputtered iridium oxide films (sirofs) for low-impedance neural stimulation and recording electrodes,” in *The 26th Annual International Conference of the IEEE Engineering in Medicine and Biology Society*, vol. 4 of *IEMBS-04*, p. 4153–4156, IEEE.

[42] S. Negi, R. Bhandari, R. Van Wagenen, and F. Solzbacher, “Factors affecting degradation of sputtered iridium oxide used for neuroprosthetic applications,” in 2010 IEEE 23rd International Conference on Micro Electro Mechanical Systems (MEMS), p. 568–571, IEEE, Jan. 2010.

[43] K. H. Chen, J. F. Dammann, J. L. Boback, F. V. Tenore, K. J. Otto, R. A. Gaunt, and S. J. Bensmaia, “The effect of chronic intracortical microstimulation on the electrode–tissue interface,” Journal of Neural Engineering, vol. 11, p. 026004, Feb. 2014.

